# Fecal microbiomes of laboratory beagles receiving antiparasitic formulations in an experimental setting

**DOI:** 10.1101/2023.08.08.552529

**Authors:** Andrea Ottesen, Brandon Kocurek, Christine Deaver, Oscar Chiesa, Rachael Cohen, Mark Mammel, Elizabeth Reed, Patrick McDermott, Errol Strain, Michael Myers

## Abstract

Here we describe the fecal microbiome of laboratory beagles in a non-invasive and humane experiment designed to contrast in vivo versus invitro bioequivalence in response to antiparasitic drug administration. The experiment provided a unique opportunity to describe the fecal microbiota of dogs in an experimental setting prior to their adoption. These data are contributed as a resource for the scientific community by the Center for Veterinary Medicine (CVM) of the U.S. Food and Drug Administration (FDA).

## Introduction

A non-invasive research study was designed to contrast in vivo pharmacokinetic values in laboratory beagles receiving formulations of Ivermectin (IVR) and Praziquantel (PZQ), with in vitro diagnostic tests [1]. Assessment of the microbiome of study dogs was not part of the original experimental design and thus, these data primarily provide a descriptive snapshot of longitudinal canine fecal microbiota in an experimental setting, pre and post exposure to formulations of IVR and PZQ. The bioequivalence study was designed and conducted under an IACUC-approved protocol using principles of humane experimental techniques, ie; Replacement, Reduction, and Refinement (RRR)[2], in compliance with the Public Health Service Policy on Humane Care and Use of Laboratory Animals (PHS Policy), and in an AAALAC-International-accredited research facility[3, 4]. These principles of animal welfare are central to FDA CVM research. All dogs were provided humane treatment and socialization throughout the study, and all dogs were adopted following completion of the study.

Research has demonstrated that many types of drugs, foods, and environments can influence the microbiome and resistome of humans and animals[5, 6]. It is well understood that states of health and disease are not only driven by host genetics, but also by the thousand-fold more abundant genetic potential of the microbiome [7, 8]. The ability to collect microbiome data pre and post exposure for human and veterinary pharmaceuticals may lead to important advancements in drug research. While primarily descriptive, these data support the hypothesis that microbiome and antimicrobial resistance changes occur as a result of exposure to antiparasitic formulations.

IVR has been used in veterinary and human medicine for more than 34 years as a broad-spectrum endo- and ectoparasitic drug [9] and also for control of onchocerciasis and lymphatic filariasis in humans[10-12]. With a half-life of more than 100 days in environmental sediments[13], this broad-spectrum biocide is likely to influence more than just its invertebrate targets and may illicit innate immune responses reflected in host microbiota. PZQ demonstrates activity against species not covered by Ivermectin. Thus, formulations of both drugs have a long history of veterinary and human use, providing expansive protection from a range of parasites[10-12].

Here, we describe the bacterial microbiota of beagles in an experimental setting before and after treatment with formulations of IVR and PZQ[1]. Metagenomic data were generated from DNA of fecal samples collected every 24 H for three days prior to treatment, three days following treatment, and time points 10, 15, 22, 109, 110, and 111 days post treatment for 4 biological replicates of 4 dog-units (2 or 3 dogs per unit) according to previously described methods[14].

Significant shifts in the relative abundance of bacterial taxa were observed 24 hours after administration of IVR and PZQ. Especially prominent were decreased abundance of *Lactobacillus* spp. and increased abundance of *Prevotella copri*. At day 10 post treatment, the relative abundance of bacterial taxa resembled pre-exposure profiles (Figure 1a). This impact is also seen in the incidence of antimicrobial resistance associated genes that are likely associated with the differentially observed bacterial taxa (Figure 1b).

**Figure 1.**
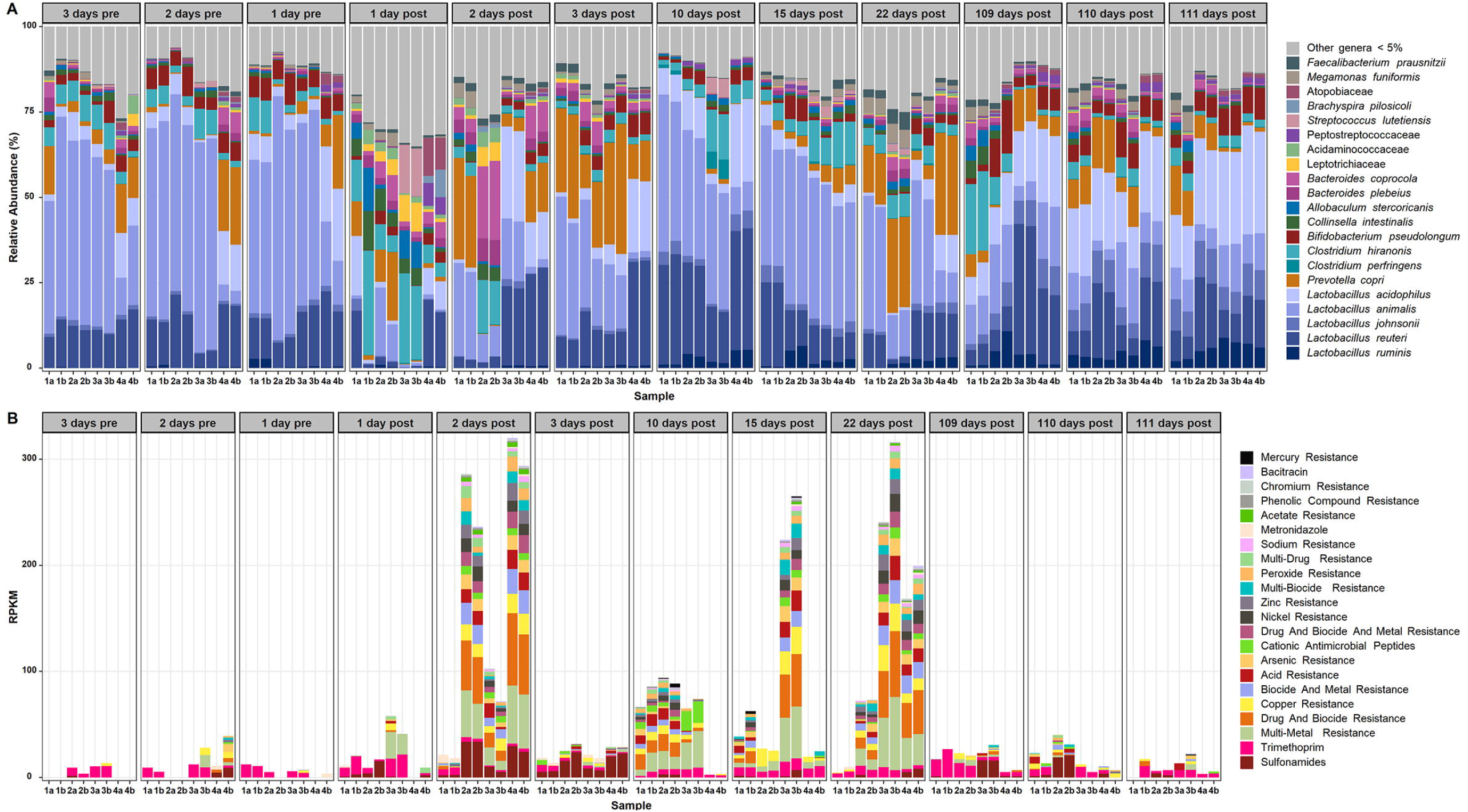
Microbiomes and resistomes of beagle feces before and after exposure to formulations of Ivermectin and Praziquantel. **Figure 1a** Microbiomes of two biological replicates from each of 4 dog-units (2-3 dogs per unit) are shown 3 days before exposure to treatment (Pre 1-3), 3 days after treatment (Post 1-3), and 10, 15, 22, 109, 110, and 111 days following treatment (Post 4-9). **Figure 1b**. The resistome (classes of genes associated with anticrobial resistance) are shown for the same biological replicates of 4 dog units shown in 1a. at the same time points pre and post treatment to IVR and PZQ.

As microbiome compositional signatures are increasingly correlated with states of health and disease and specific features are definitively identified for use as diagnostics, an exciting new frontier of high-resolution microbiome-based diagnostics and treatment is pushing forward [15, 16]. Implicit in this frontier is the need for a new element of due diligence in drug evaluation which includes assessment of the impact of drug formulations on host microbiomes.

## Data Availability

Sequence data for 4 biological replicates of 4 dog units at 12 timepoints across 111 days of study have been deposited at the National Center for Biological Information (NCBI) under the canine metagenome Bioproject (PRJNA1003522) with biosample accession numbers SAMN36892673 through SAMN36892768. The data was deposited at the NCBI using the MIMS Environmental/Metagenome host associated package.

## Notes

### Competing Interest Statement

The authors have declared no competing interest.

